# The intrinsically disordered C-terminal linker of FtsZ regulates protofilament dynamics and superstructure *in vitro*

**DOI:** 10.1101/171280

**Authors:** Kousik Sundararajan, Erin D. Goley

**Affiliations:** From the Department of Biological Chemistry, Johns Hopkins University School of Medicine, Baltimore, Maryland, 21205, United States of America

**Keywords:** FtsZ, *Caulobacter crescentus*, intrinsically disordered, bundling, cytoskeleton

## Abstract

The bacterial tubulin FtsZ^2^ polymerizes to form a discontinuous cytokinetic ring that drives bacterial cell division by directing local cell wall synthesis. FtsZ comprises a polymerizing GTPase domain, an intrinsically disordered C-terminal linker (CTL) and a C-terminal conserved α-helix (CTC). FtsZ protofilaments align circumferentially in the cell, with the CTC mediating attachment to membrane-associated division proteins. The dynamic turnover and treadmilling of clusters of FtsZ protofilaments guides cell wall synthesis and constriction. The nature and regulation of the interactions that result in the assembly of protofilaments into dynamic clusters is unknown. Here, we describe a role for the CTL of *Caulobacter crescentus* FtsZ as an intrinsic regulator of lateral interactions between protofilaments *in vitro*. FtsZ lacking its CTL (ΔCTL) shows dramatically increased propensity to form long multifilament bundles compared to wildtype (WT). ΔCTL has reduced GTP hydrolysis rate compared to WT. However, reducing protofilament turnover in WT is not sufficient to induce bundling. Surprisingly, binding of the membrane-anchoring protein FzlC disrupts ΔCTL bundling in a CTC-dependent manner. Moreover, the CTL affects the ability of FtsZ curving protein FzlA to promote formation of helical bundles. We conclude that the CTL of FtsZ influences polymer structure and dynamics both through intrinsic effects on lateral interactions and turnover and by influencing extrinsic regulation of FtsZ by binding partners. Our characterization of CTL function provides a biochemical handle for understanding the relationship between Z-ring structure and function in bacterial cytokinesis.

---

Canonical cytoskeletal proteins in animal cells polymerize to form structural elements that provide shape and mechanical integrity to the cell. In bacteria, however, the cell wall is the primary structural element, maintaining cell shape and preventing lysis. The role of bacterial cytoskeletal proteins that impact cell shape is in the spatial and temporal regulation of cell wall synthesis (1, 2). Cytoskeletal polymer assembly, structure, and dynamics collectively regulate local shape changes by constraining and/or directing cell wall remodeling enzymes (2). During cell division, the cytoskeletal protein FtsZ polymerizes to form the cytokinetic ring or “Z-ring” at the incipient division site and recruits over two dozen proteins, including cell wall enzymes (3–5). The Z-ring comprises discontinuous clusters of circumferentially aligned FtsZ protofilaments that are highly dynamic (6–9). How protofilaments are arranged within the resolution-limited clusters and how they interact with each other is largely unknown.

Recent studies in *Escherichia coli* and *Bacillus subtilis* demonstrated that FtsZ protofilament clusters in the Z-ring undergo treadmilling motion (8, 9). FtsZ treadmilling, in turn, drives the circumferential movement of cell wall enzymes, thereby directing local cell wall synthesis and remodeling towards constriction. The treadmilling of FtsZ clusters is presumably driven by the polymerization and depolymerization of FtsZ protofilaments, since GTP hydrolysis mutants that have reduced turnover *in vitro* lead to slower movement of clusters *in vivo* (8, 9). In addition to turnover of polymers, other aspects of the Z-ring such as conformational changes within FtsZ, lateral interactions between protofilaments and interaction with FtsZ-binding proteins likely regulate cluster movement and thereby, local cell wall remodeling (4, 10, 11). For example, FtsZ binding proteins such as ZapA in *E. coli* and FzlA in *C. crescentus* that cause cross-linking or bundling of FtsZ protofilaments *in vitro* are important for efficient cytokinesis through unclear mechanisms (12–14).

FtsZ has a tubulin-like GTPase domain that polymerizes on binding GTP (15–17) (Figure 1A). GTP hydrolysis is stimulated by polymerization, and destabilizes the monomer-monomer interface, leading to depolymerization (17, 18). Under physiologically relevant conditions and in the presence of GTP, FtsZ predominantly forms straight or gently curved single protofilaments and/or double protofilament bundles *in vitro* by transmission electron microscopy (19, 20). In cells, FtsZ protofilaments are recruited to the membrane by membrane anchoring proteins that bind FtsZ through the conserved α-helix at its extreme C-terminus (CTC or C-terminal conserved peptide) (3, 21–24). Between the GTPase domain and the CTC is an unstructured region called the C-terminal linker or CTL that varies widely in length (2 – 330 amino acids) and sequence across species (21). FtsZ from most bacteria with a peptidoglycan cell wall that have a CTC have a minimum CTL of 9 amino acids (21). Studies in *E. coli* and *B. subtilis* suggest length-, flexibility-, and/or disorder-dependent roles for the intrinsically disordered CTL in determining FtsZ assembly and function *in vivo* (25–27).

**FIGURE 1:**
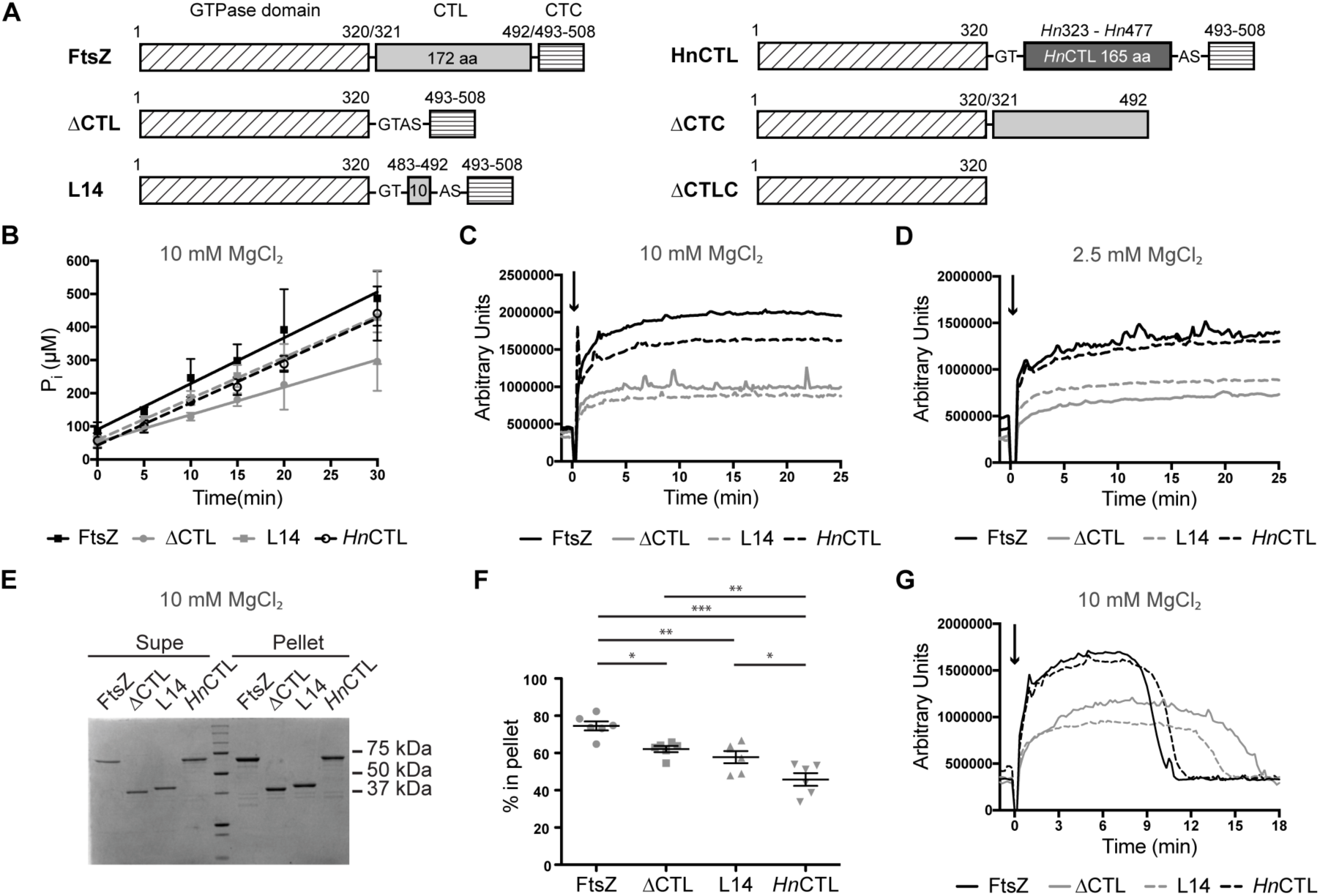
CTL affects FtsZ polymer dynamics **A.** Schematic showing different regions of FtsZ, truncations and CTL variants. The numbers shown above schematic correspond to the amino acid numbers in wild type (*Cc*FtsZ). *Hn*323 - *Hn*477 represent amino acid numbers in *Hyphomonas neptunium* FtsZ. (G, T, A and S represent amino acids glycine, threonine, alanine and serine introduced when restriction sites were added for cloning.) **B.** Inorganic phosphate (P_i_) concentration over time in reactions containing 4 μM FtsZ CTL variants with 2 mM GTP and 10 mM MgCl_2_ (n=3). Error bars represent standard deviation. Straight lines indicate linear fits of averages. **C.** Right angle light scatter at 350 nm over time for 4 μM FtsZ CTL variants with 2 mM GTP (added at time = 0 min) and 10 mM MgCl_2_ (mean of 3 replicates). **D.** Right angle light scatter at 350 nm over time for 4 μM FtsZ CTL variants with 2 mM GTP (added at time = 0 min) and 2.5 mM MgCl_2_ (mean of at least 3 replicates). **E.** A representative Coomassie stained SDS-PAGE gel of a pelleting assay showing relative amounts of FtsZ CTL variants in supernatant (Supe) or pellet for 4 μM FtsZ CTL variant with 2 mM GTP and 10 mM MgCl_2_, 15 minutes after addition of GTP. **F.** Quantification of the percentage of FtsZ CTL variants in pellet corresponding to experiment in E (n = 6). * - p < 0.05, ** - p < 0.01, *** - p < 0.001 for non-parametric t-tests. **G.** Right angle light scatter at 350 nm over time for 4 μM FtsZ CTL with 0.5 mM GTP (added at time = 0 min) and 10 mM MgCl_2_.

In our previous study characterizing CTL function in *Caulobacter crescentus*, we observed that *C. crescentus* FtsZ lacking its 172 amino-acid long CTL (ΔCTL) forms highly bundled protofilaments *in vitro* (28) (Figure 1A). This mutant of FtsZ when expressed *in vivo* leads to filamentation due to cytokinesis failure but also causes dominant lethal local cell wall defects leading to cell envelope bulges and rapid lysis. We were surprised to find that a variant of FtsZ with a 14-amino acid CTL (L14, Figure 1A) caused filamentation but no bulging or lysis. Moreover, the L14 variant formed less bundled structures *in vitro* (28). We hypothesized that the aberrant assembly properties of ΔCTL *in vitro* underlie the defects in the structure and function of the Z-ring formed by ΔCTL *in vivo* (28).

In the present study, we sought to fully characterize the contributions of the CTL to the polymerization properties of FtsZ *in vitro.* To this end, we compared polymers formed by wildtype FtsZ with ΔCTL and L14. We also included in our *in vitro* characterization a chimeric FtsZ variant with the *Hyphomonas neptunium* CTL in place of the *C. crescentus* CTL (*Hn*CTL) (Figure 1A). *Hn*CTL is inefficient but functional for cytokinesis – while the Hn*CTL* gene can replace *ftsZ* at its genomic locus in *C. crescentus*, the resulting cells have slower doubling time and heterogeneous cell length (28). Our *in vitro* characterization of the CTL variants of FtsZ and their interactions with FtsZ binding factors has revealed that the CTL is important for protofilament turnover and lateral interaction. ΔCTL tends to form long extended bundles that require GTP binding but not hydrolysis for their formation. The membrane-anchoring protein FzlC can disrupt bundle formation in a CTC-dependent manner, but the CTC itself does not contribute to bundling. Moreover, the FtsZ curving protein FzlA can no longer robustly form helical bundles with ΔCTL. Overall, our study provides a biochemical framework to connect the assembly properties of these CTL variants *in vitro* with their *in vivo* Z-ring structures, dynamics, and effects on the regulation of cell wall modeling.

## RESULTS

### FtsZ-CTL contributes to polymer dynamics

To better understand the contributions of the CTL to FtsZ polymerization, we performed a series of biochemical assays using purified CTL variants – FtsZ (WT), ΔCTL, L14 and *Hn*CTL – *in vitro.* Since polymerization and depolymerization are coupled to GTP binding and hydrolysis, we first determined the GTP hydrolysis rate for each variant using a malachite green assay to detect inorganic phosphate liberated over time. We observed a reduction in the rate of GTP hydrolysis for each CTL variant as compared to WT (Figure 1B), with ΔCTL exhibiting the slowest rate (Figure 1B) (28). At 10 mM MgCl_2_ and 4 μM WT or CTL variants, WT had a GTP hydrolysis rate of 3.46 ± 0.37 min^−1^, while *Hn*CTL, L14 and ΔCTL had rates of 3.20 ± 0.21, 3.11 ± 0.33 and 2.07 ± 0.51 min^−1^ respectively.

To monitor polymerization over time for the FtsZ CTL variants, we used right angle light scattering. Light scattering has been used as a read out of FtsZ polymerization, as polymers scatter more light than monomers and the amount of light scatter increases with the size of polymers (19, 20). It is important to keep in mind, however, that light scattering is also influenced by monomer size and structure and by formation of higher order filament assemblies such as bundles. Using this method for reactions containing 10 mM MgCl_2_ and 2 mM GTP, we found that all variants readily polymerized upon addition of GTP with no detectable lag phase at our time resolution, and we did not observe any obvious differences in their initial rates of increase in scatter (Figure 1C). However, whereas WT and *Hn*CTL resulted in similar light scatter values at steady state, steady state light scatter values for ΔCTL and L14 were significantly lower (Figure 1C). This trend was also true at the presumed physiological MgCl_2_ concentration of 2.5 mM. (Figure 1D).

Curiously, when we determined the extent of polymerization of the FtsZ CTL variants using high speed pelleting in the presence of 10 mM MgCl_2_, we were surprised to observe a distinct trend for steady state polymer mass. While each CTL variant showed a reduced percentage in the pellet (representing polymer fraction) compared to WT, ΔCTL and L14 were enriched in the pellet compared to *Hn*CTL (Figure 1E, 1F). We reason that the longer CTLs of WT and *Hn*CTL may significantly contribute to the light scattering properties of their resulting polymers and that formation of higher order structures by one or more of the CTL variants may also influence light scattering or propensity to pellet, leading to different results in these two assays.

Finally, we used limiting concentrations of GTP to monitor the decrease in light scatter as filaments depolymerize following consumption of GTP. We found that the FtsZ CTL variants with slower GTP hydrolysis rates took longer to return to baseline, indicating a strong correlation between the effect of the CTL on GTP hydrolysis rates and polymer dynamics, as we would predict (Figure 1G).

### The CTL influences lateral interaction between protofilaments

We next assessed the filament structures formed by the four CTL variants using negative stain transmission electron microscopy (TEM). We observed polymers with similar protofilament widths for the four variants, but with distinct appearance and abundance depending on the nature of the CTL. Similar to previous reports for *C. crescentus* FtsZ (12, 20, 29, 30), at 8 μM protein concentration with GTP and 2.5 mM MgCl_2_ WT FtsZ formed gently curved single filaments and occasional straight double filament bundles (Figure 2A). *Hn*CTL and L14 predominantly formed sparse, short single filaments under the same conditions. Strikingly, as reported earlier (28), ΔCTL predominantly forms long multi-filament bundles at 8 μM protein concentration and 2.5 mM MgCl_2_. We did not observe these long bundled structures for the other variants (Figure 2A).

**FIGURE 2:**
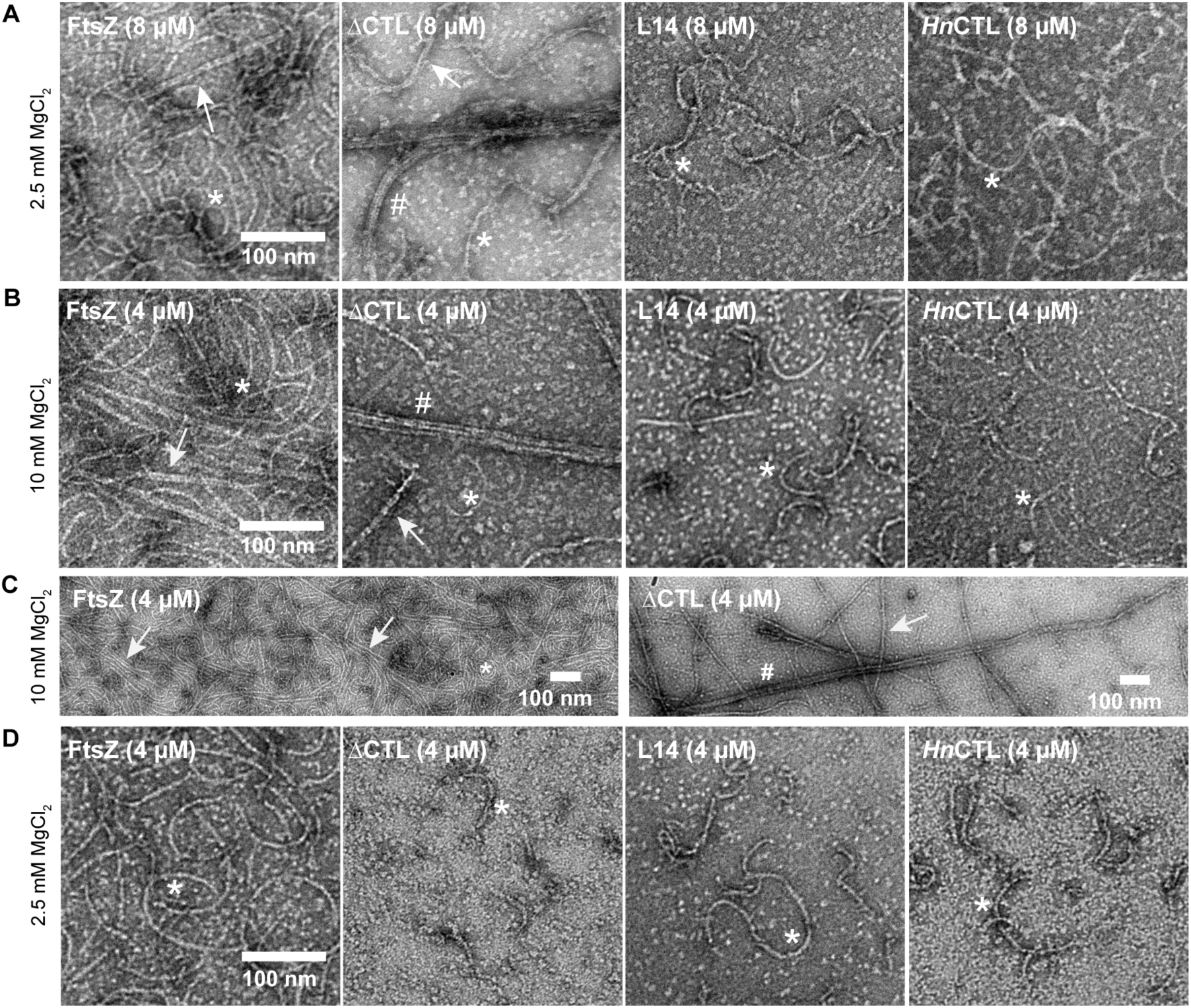
CTL affects lateral interaction between FtsZ protofilaments. Electron micrographs of polymers formed by FtsZ CTL variants spotted on grids 15 minutes after addition of GTP and stained with uranyl formate. **A.** 8 μM FtsZ CTL variant with 2 mM GTP and 2.5 mM MgCl_2_. **B.** 4 μM FtsZ CTL variant with 2 mM GTP and 10 mM MgCl_2_. **C.** Low magnification micrographs of FtsZ or ΔCTL polymers with 2 mM GTP and 10 mM MgCl_2_. **D.** 4 μM FtsZ CTL variant with 2 mM GTP and 2.5 mM MgCl_2_. Scale bars – 100 nm. * - single protofilaments, arrow – two or three filament bundles, # - multifilament bundles.

Increasing the concentration of MgCl_2_ has been reported to promote lateral interactions between FtsZ protofilaments. We therefore explored the effect of high MgCl_2_ concentration on polymer superstructure for the four FtsZ variants. WT FtsZ frequently formed two- or three-filament bundles (7 - 15 nm wide) at 10 mM MgCl_2_ (Figure 2B, 2C). In contrast, under the same conditions ΔCTL bundles often consisted of more than 4 filaments (~20 - 200 nm in width) and appeared as very long, straight structures frequently extending longer than 1 micron (Figure 2B, 2C). At 4 μM protein and 2.5 mM MgCl_2_ concentration, we did not observe bundles for ΔCTL (Figure 2D). Compared to WT protofilaments, L14 and *Hn*CTL protofilaments were sparse on the grids and were predominantly gently-curved single filaments at either MgCl_2_ concentration (Figure 2B, 2D).

We further investigated if polymer stability and/or slow GTP hydrolysis contributed to the increased bundling observed for ΔCTL *in vitro.* To test this, we used GMPCPP – a slowly hydrolyzed analog of GTP – to form stable protofilaments at high and low concentrations of MgCl_2_. By TEM, at 2.5 mM MgCl_2_ concentration, the polymer structures were not significantly different with GMPCPP compared to GTP (Figure 3A). At 10 mM MgCl_2_, we readily observed large multifilament bundles of ΔCTL protofilaments with GMPCPP similar to those observed with GTP (Figure 3B). While we observed more protofilaments for all the FtsZ CTL variants, and more interaction between protofilaments for WT, we failed to find the long, wide multi-filament bundles characteristic of ΔCTL for the other variants. We conclude that the ability to form long multifilament bundles is specific to ΔCTL.

**FIGURE 3:**
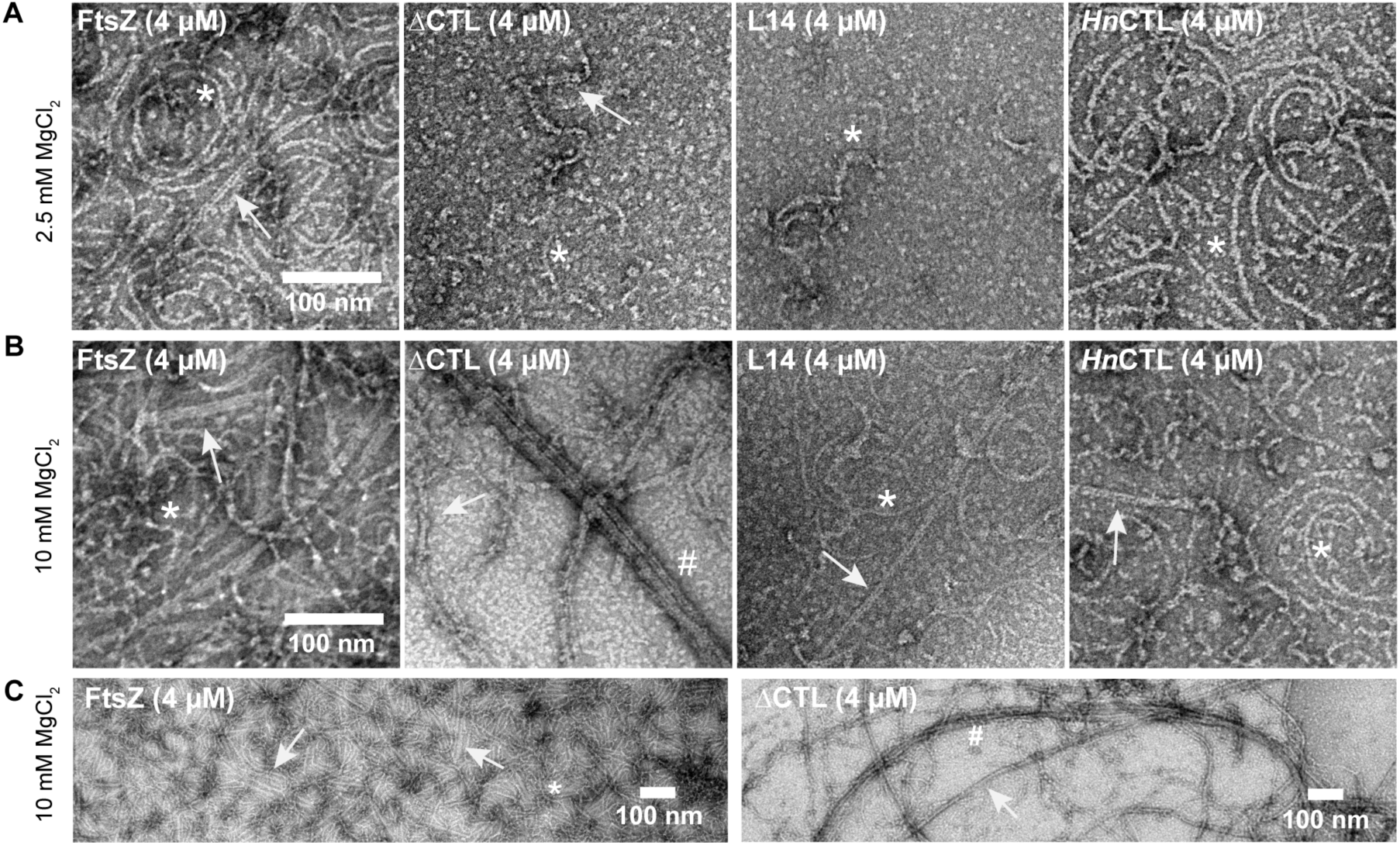
Effects of CTL on protofilament lateral interaction are independent of GTP hydrolysis. Electron micrographs of polymers formed by FtsZ CTL variants spotted on grids 15 minutes after addition of GMPCPP and stained with uranyl formate. **A.** 4 μM FtsZ CTL variant with 0.2 mM GMPCPP and 2.5 mM MgCl_2_. **B.** 4 μM FtsZ CTL variant with 0.2 mM GMPCPP and 2.5 mM MgCl_2_. **C.** Low magnification micrographs of FtsZ or ΔCTL polymers with 0.2 mM GMPCPP and 10 mM MgCl_2_. Scale bars – 100 nm. * - single protofilaments, arrow – two or three filament bundles, # - multifilament bundles.

The influence of CTL on lateral interaction was also evident from the differences in light scatter profiles of the CTL variants with GMPCPP (Figure 4). At 2.5 mM MgCl_2_, we did not observe large differences in the kinetics or extent of light scatter over time with GTP or GMPCPP for the CTL variants (Figure 4A, 4B). Only WT showed significantly higher scatter with GMPCPP compared to GTP (Figure 4B). However, at 10 mM MgCl_2_ with GMPCPP, we observed a two-step increase in light scatter for WT and ΔCTL (Figure 4C). For WT we observed an initial rapid increase immediately following the addition of nucleotide (similar to light scatter with GTP) and then a second slower, much larger increase about 5 minutes later – likely corresponding to polymerization into protofilaments and formation of bundles respectively. For ΔCTL, the initial rapid increase was almost immediately followed by the second increase (after about 2 minutes). Moreover, the secondary increase in light scatter was much faster and larger for ΔCTL compared to WT (Figure 4C, 4D). *Hn*CTL and L14 only undergo a rapid one-step increase in light scatter after addition of GMPCPP, not obviously different from GTP (Figure 4C, 4D) likely due to the absence of higher order assemblies of protofilaments under these conditions.

**FIGURE 4:**
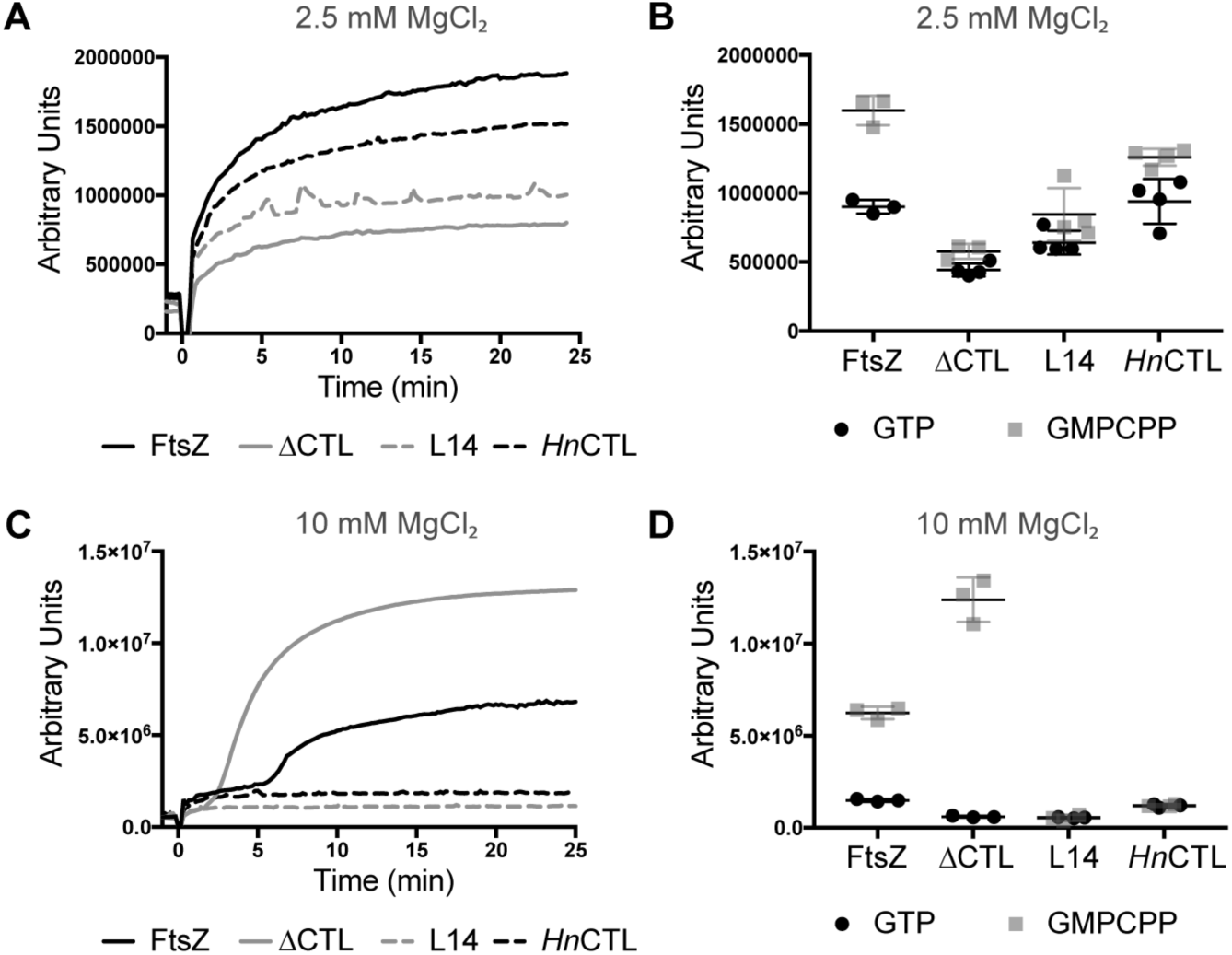
Polymerization kinetics of CTL variants with GMPCPP. **A.** Right angle light scatter at 350 nm over time for 4 μM FtsZ CTL variants with 0.2 mM GMPCPP (added at time = 0 min) and 2.5 mM MgCl_2_ (mean of 3 replicates). **B.** Extent of light scatter (quantified as the difference between light scatter at steady state (time = 25 min) and light scatter prior to addition of nucleotide (time = −1 min)) for FtsZ CTL variants with GTP (from Figure 1D) or GMPCPP (Figure 4A). **C.** Right angle light scatter at 350 nm over time for 4 μM FtsZ CTL variants with 0.2 mM GMPCPP (added at time = 0 min) and 10 mM MgCl_2_ (mean of 3 replicates). **D.** Extent of light scatter (quantified as the difference between light scatter at steady state (time = 25 min) and light scatter prior to addition of nucleotide (time = −1 min)) for FtsZ CTL variants with GTP (from Figure 1C) or GMPCPP (Figure 4C).

We confirmed that the second increase in light scatter for WT and ΔCTL with GMPCPP at 10 mM MgCl_2_ were indeed due to increased lateral interaction by observing polymer structure over time (Figure 5). WT incubated with GTP and 10 mM MgCl_2_ forms predominantly short single filaments after 30 seconds and both single filaments and double or triple filament bundles after 15 minutes (Figure 5A). In contrast, ΔCTL incubated with GTP and 10 mM MgCl_2_ forms few straight bundles that become longer and thicker with time (Figure 5B). The difference in polymer structure for WT and ΔCTL over time was more apparent with GMPCPP and 10 mM MgCl_2_ (Figure 5C, 5D). While WT largely formed single gently curved protofilaments after incubation with GMPCPP for 30 seconds, we observed more double- and triple-filament bundles after 5 minutes or longer incubation (Figure 5C). We also observed a few long four or five filament bundles for WT after incubation with GMPCPP for 25 minutes. In contrast, we almost exclusively observe long four- or five-filament bundles for ΔCTL after incubation with GMPCPP for 30 seconds (Figure 5D). ΔCTL multifilament bundles were more prominent, longer and thicker after longer incubation with GMPCPP. Taking our observations together, we infer that the CTL is important for reducing lateral interaction between FtsZ protofilaments and that a minimal CTL of 14 amino acids is sufficient to perform this role.

**FIGURE 5:**
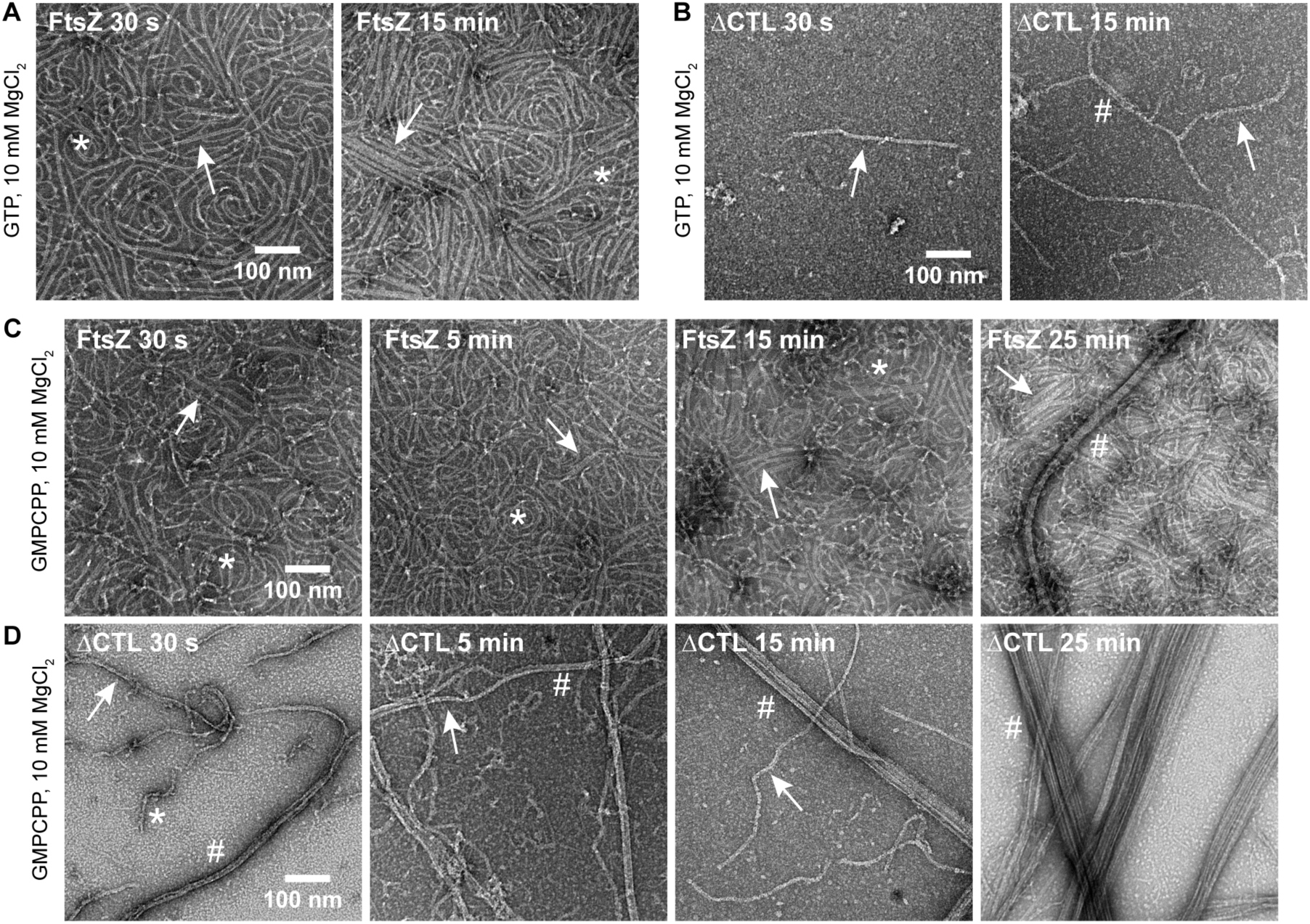
ΔCTL becomes more bundled with time. Electron micrographs of polymers formed by FtsZ or ΔCTL spotted on grids at the indicated amount of time after addition of nucleotide and stained with uranyl formate. **A.** 4 μM FtsZ with 2 mM GTP and 10 mM MgCl_2_. **B.** 4 μM ΔCTL with 2 mM GTP and 10 mM MgCl_2_. **C.** 4 μM FtsZ with 0.2 mM GMPCPP and 10 mM MgCl_2_. **D.** 4 μM ΔCTL with 0.2 mM GMPCPP and 10 mM MgCl_2_. Scale bars – 100 nm. * - single protofilaments, arrow – two or three filament bundles, # – multifilament bundles

Curiously, *Hn*CTL does not form prominent double- or triple-filament bundles under any condition tested despite having WT-like light scatter with GTP. The decreased ability of *Hn*CTL to form higher order protofilament assemblies is further suggested by the lower percentage of *Hn*CTL in the polymer fraction as observed by high-speed pelleting (Figure 1E, 1F). A technical explanation for the sparse, unbundled filaments observed for *Hn*CTL by TEM might be that *Hn*CTL does not stick to the grids for TEM as well as WT FtsZ. However, with GMPCPP it is clear from light scattering that WT FtsZ forms higher order structures while *Hn*CTL does not. Thus, the sequence of the CTL seems to play a role in maintaining optimal interfilament interactions in a way that is altered for *Hn*CTL.

### High MgCl_2_-concentration does not increase polymerization of ΔCTL

We observe bundles of ΔCTL with GTP or GMPCPP more readily at 10 mM MgCl_2_ compared to 2.5 mM MgCl_2_. This effect of MgCl_2_ on increasing lateral interaction has been documented before for full length *E. coli* FtsZ, with more two- or three-filament bundles at higher MgCl_2_ concentrations (19). We see the same effect for *C. crescentus* FtsZ (WT) (compare TEM shown in Figures 2B and 2D or Figures 3A and 3B). However, we see fewer protofilaments for ΔCTL by EM and lower signal by light scattering at 2.5 mM MgCl_2_ than at 10 mM MgCl_2_. We reasoned that the effects of increased divalent cation concentration might be the result of increased polymer formation (i.e. increase in total protofilament mass), increased bundling, or both. To differentiate between these possibilities, we used the polymerization-specific fluorescence reporter - a tryptophan mutant of FtsZ (FtsZ_L72W_) developed by the Erickson lab (29). This mutant has a tryptophan residue engineered on the polymerizing interface of FtsZ, the environment of which changes on polymerization resulting in a polymerization-specific increase in fluorescence signal. Unlike light scattering, the increase in tryptophan fluorescence on polymerization is minimally affected by bundling and is a more reliable readout of number of the longitudinal interactions.

Using this assay, we first added GMPCPP to reactions containing 4 μM FtsZ (with 10% FtsZ_L72W_) or ΔCTL (with 10% ΔCTL_L72W_) then initiated formation of polymers with 2.5 mM MgCl_2_ (Figure 6A). We observed a polymerization-dependent increase in tryptophan fluorescence for both WT and ΔCTL, as expected. Then we added MgCl_2_ up to 10 mM and continued to measure the fluorescence intensity. Since the excitation and emission wavelengths for right angle light scatter and tryptophan fluorescence are similar (350 nm – 350 nm compared to 295 nm – 344 nm respectively), we also included FtsZ alone or ΔCTL alone (with no tryptophan mutation) as controls for a possible increase in intensity readout as a result of scatter. For WT, with or without the tryptophan mutant, we failed to observe a change in intensity when MgCl_2_ concentration was increased, suggesting that increased MgCl_2_ does not change the polymer mass at steady state under these conditions. For ΔCTL, we did observe an increase in signal when we increased the MgCl_2_ concentration. However, this signal increase was observed for the ΔCTL alone control, as well, indicating that it was the result of bundling-induced light scatter. Indeed, when we subtract the fluorescence signal observed for the ΔCTL alone control from that of ΔCTL with the tryptophan mutant, we observe no additional increase in signal on increasing MgCl_2_ concentration (Figure 6B). These observations argue against the possibility that the increased bundling seen by EM and increased signal observed by light scattering at high MgCl_2_ concentration is due to increased protofilament mass.

**FIGURE 6:**
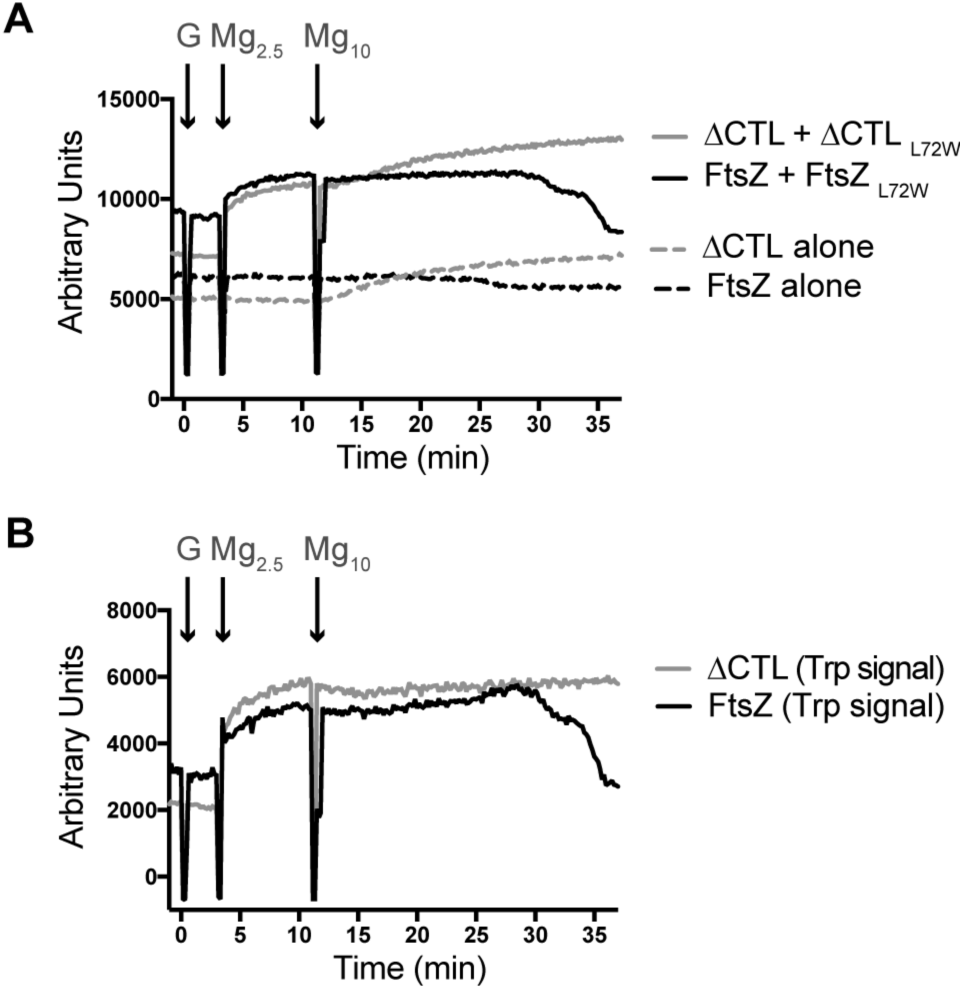
Effect of MgCl_2_ on polymerization of FtsZ or ΔCTL. **A.** Fluorescence signal emitted over time (emission at 344 nm after excitation at 295 nm) for FtsZ or ΔCTL with 0.1 mM GMPCPP added at time = 0 min. MgCl_2_ was added to reaction to a concentration of 2.5 mM at time = 3 min and subsequently to a concentration of 10 mM at time = 11 min (mean of 3 replicates). Total concentration of protein in each reaction was 4 μM. L72W mutants when used were at 10% total protein concentration (3.6 μM FtsZ + 0.4 μM FtsZ_L72W_, 3.6 μM ΔCTL + 0.4 μM ΔCTL_L72W_, 4 μM FtsZ alone or 4 ΔCTL alone). **B.** Signal specific to tryptophan fluorescence (calculated by subtracting mean of signal for FtsZ or ΔCTL alone from mean of corresponding signal for FtsZ + FtsZ_L72W_ or ΔCTL + ΔCTL_L72W_ respectively) for experiment in 6A.

We also observed a drop in tryptophan fluorescence signal back to baseline after some time for WT, potentially due to the eventual depletion of GMPCPP through slow hydrolysis (Figure 6A, 6B). In the same time frame, we did not observe a similar decrease for ΔCTL likely because it has a slow nucleotide hydrolysis rate compared to WT.

### FtsZ and ΔCTL can copolymerize in vitro

Since ΔCTL has a dominant effect on cell shape and division in cells, we hypothesized that ΔCTL and WT FtsZ co-polymerize *in vivo* and that ΔCTL exerts its propensity for increased lateral interaction in the resulting copolymers. Since the polymerizing GTPase domain is intact in ΔCTL, we expected that WT and ΔCTL could still copolymerize. To test this formally, first, we empirically determined the critical concentrations for polymerization of the tryptophan mutants FtsZ_L72W_ and ΔCTLL72W both by GTP hydrolysis measurements and by determining the concentration below which no increase in tryptophan fluorescence was observed on addition of GTP. Then, we used the tryptophan mutants of FtsZ or ΔCTL at concentrations below their respective critical concentrations along with ΔCTL or FtsZ to assay copolymerization. Using only FtsZ_L72W_ below its critical concentration for polymerization, we observed no increase in fluorescence on addition of GTP at 2.5 mM MgCl_2_ concentration (Figure 7A). When we added WT FtsZ to this reaction mixture, pushing the total protein concentration above the critical concentration for FtsZ polymerization, we observed a significant GTP-dependent increase in fluorescence (Figure 7A). We inferred from this observation that WT FtsZ can copolymerize with FtsZ_L72W_ and pushes the overall monomer concentration above the critical concentration, thereby resulting in polymerization and an increase in tryptophan fluorescence. Importantly, we also observed this fluorescence increase when we added ΔCTL to the reaction mixture instead of FtsZ, indicating that FtsZ_L72W_ could copolymerize with ΔCTL as well as WT. We observed the same effect when we used ΔCTL_L72W_ instead of FtsZ_L72_w further confirming that FtsZ and ΔCTL can copolymerize (Figure 7B). Since we used limiting concentrations of GTP (50 μM), we could also observe the decrease in fluorescence intensity due to depolymerization following depletion of GTP. Time taken until depolymerization was exactly as estimated from the individual GTP hydrolysis rates, dominated by the predominant species of monomers (FtsZ or ΔCTL), further adding confidence to our results. We did not observe any significant increase in fluorescence signal at 344 nm on addition of GTP for 4 μM FtsZ or ΔCTL at 2.5 mM MgCl_2_ (Figure 7C), suggesting that the signal observed in the presence of tryptophan mutants are indicative of polymerization and not background scatter from polymerization of the proteins lacking the tryptophan.

**FIGURE 7:**
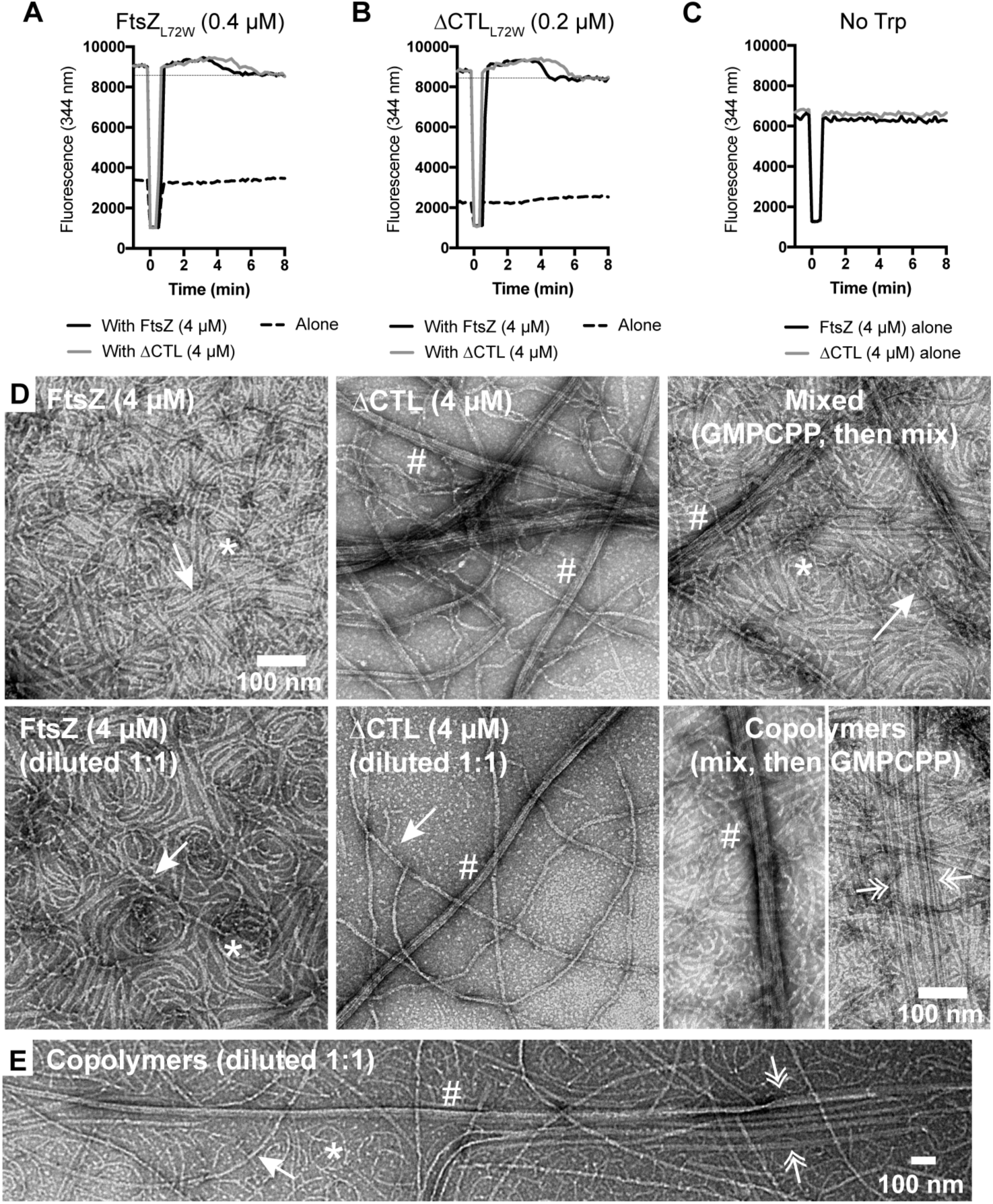
FtsZ and ΔCTL can copolymerize to form very long bundles. **A - C.** Tryptophan fluorescence increase (emission at 344 nm after excitation at 295 nm) for FtsZ CTL variants with 50 μM GTP added at time = 0 min and 2.5 mM MgCl_2_. Straight black line above 8000 indicating baseline after depolymerization has been included to better visualize the increase in fluorescence after addition of GTP. **A.** Fluorescence signal from 0.4 μM FtsZ_L72W_ in reaction with 4 μM FtsZ or ΔCTL or by itself (n=3). **B.** Fluorescence signal from 0.2 μM ΔCTL_L72W_ in reaction with 4 μM FtsZ or ΔCTL or by itself (n=3). **C.** Fluorescence signal from 4 μM FtsZ or ΔCTL (n=1). **D.** Electron micrographs showing polymers formed by FtsZ or ΔCTL individually, mixed together (after polymerization individually) or FtsZ:ΔCTL copolymers with 0.2 mM GMPCPP and 10 mM MgCl_2_ spotted on EM grids 15 minutes after addition of GMPCPP and stained with uranyl formate. Mixed represents micrographs of 4 μM FtsZ and 4 μM ΔCTL polymerized individually and then mixed prior to spotting on grids. FtsZ (diluted to 1:1) or ΔCTL (diluted 1:1) represent polymers obtained by diluting GMPCPP-stabilized polymers of 4 μM FtsZ or ΔCTL respectively with polymerization buffer (1:1 dilution) prior to spotting on grids. Copolymer represents micrographs of polymers formed by FtsZ (2 μM) and ΔCTL (2 μM) both in the same reaction mixture (total FtsZ concentration - 4 μM) with GMPCPP. **E.** Low magnification micrographs of copolymers formed by 2 μM FtsZ and 2 μM ΔCTL (initial total FtsZ concentration – 4 μM), diluted 1:1 with polymerization buffer (final total FtsZ variant concentration – 2 μM) following 15 minute incubation with 0.2 mM GMPCPP and 10 mM MgCl_2_ prior to spotting on grids. Dilution of GMPCPP-stabilized polymers reduces crowding on EM grids and helps in better visualization of polymer structures. Scale bar – 100 nm. * - single protofilaments, arrow – two or three filament bundles, # – multifilament bundles, pair of double arrowheads – groups of two- or three-filament bundles that run parallel.

On confirming that FtsZ and ΔCTL can copolymerize, we compared the polymer structures formed by the copolymers at different ratios of FtsZ to ΔCTL using negative stain TEM (Supplementary figure 1). Specifically, we compared the structures observed for mixtures of ΔCTL and WT FtsZ with those formed at the same concentrations of WT FtsZ alone at 10 mM MgCl_2_ concentration with GTP. We observed more polymer density on the EM grids with increasing FtsZ:ΔCTL ratios (Supplementary figure 1A). We also observed more bundles for FtsZ:ΔCTL combinations compared to WT alone (compare Supplementary figure 1A and first panel of 1B). We saw prominent long ΔCTL-like bundles at the lowest FtsZ:ΔCTL ratio (1:9). At higher ratios (1:3 or 1:1), the polymers were predominantly double or triple filament bundles similar to WT. However, the very tight packing of WT FtsZ on the grids made it difficult to distinguish possible differences in the length and abundance of bundles formed by FtsZ alone vs copolymers. The slowed polymer turnover of ΔCTL could potentially lead to a higher ratio of FtsZ:ΔCTL in the polymer at steady state compared to initial conditions, further complicating the interpretation of these observations by TEM.

To more clearly visualize whether a 1:1 copolymer of FtsZ:ΔCTL exhibits different filament architecture compared to individual polymers of FtsZ or ΔCTL, we compared GMPCPP-stabilized protofilaments formed with a mixture of 2 μM FtsZ and 2 μM ΔCTL to those formed by 2 μM FtsZ or 2 μM ΔCTL alone (Figure 7D). The use of GMPCPP-stabilized filaments allowed us to also visualize a 1:1 mixture of separately pre-polymerized protofilaments of FtsZ and ΔCTL to better identify structures unique to copolymers. We observed polymer structures characteristic of both FtsZ (predominantly gently curved single filaments or two- or three-filament bundles) and ΔCTL (long multifilament bundles) on the grids with a mixture of pre-polymerized protofilaments (Figure 7D, Mixed (2 μM each)). FtsZ:ΔCTL copolymers had a range of polymer structures from individual curved protofilaments to long double filament bundles to large multifilament bundles (Figure 7D, Copolymers (4 μM), Figure 7E). Uniquely, the FtsZ:ΔCTL copolymers also formed very long, evenly spaced double protofilament bundles that were not seen for mixed or individual polymers of FtsZ and/or ΔCTL (Figure 7D, 7E, double arrow). These long double filament bundles were conspicuous, as they often were observed in evenly spaced parallel arrays on the grid.

Overall, our results confirm that FtsZ and ΔCTL can copolymerize and that FtsZ:ΔCTL copolymers can have distinct structural properties, most notably propensity to bundle and length of polymers, depending on the ratio of FtsZ:ΔCTL. While it is not clear if ΔCTL’S propensity to form multi-filament bundles is relevant *in vivo*, these observations are in agreement with the increased time taken to elicit the ΔCTL bulging and lysis phenotype in the presence of WT FtsZ compared to in its absence (28).

It is also interesting to note that we still see long thick ΔCTL bundles on EM grids on dilution after polymerization with GMPCPP (compare ΔCTL (4μM) to ΔCTL (diluted 1:1) in Figure 7D), whereas WT FtsZ double or triple filament bundles were less obvious on dilution after polymerization with GMPCPP (compare FtsZ (4μM) to FtsZ (diluted 1:1) in Figure 7D). This further suggests that the increased ΔCTL interaction is not an artefact of crowding on the EM grids but an ability to form stable lateral interactions that is lacking in WT FtsZ.

### Increased lateral interaction between ΔCTL protofilaments is CTC-independent

We previously reported that ΔCTL requires its extreme C-terminus (C-terminal Conserved helix or CTC) to cause cell wall defects (28). *In vivo*, ΔCTLC (FtsZ lacking both the CTL and CTC, Figure 1A) forms elongated structures in the cytoplasm instead of Z-rings (28). In contrast, ΔCTC (FtsZ lacking only its CTC, Figure 1A) forms broad Z-rings. To test if the CTC contributes to the increased lateral interaction between ΔCTL protofilaments *in vitro*, we compared protofilaments formed by ΔCTC and ΔCTLC to ΔCTL bundles *in vitro* by TEM. At low MgCl_2_ concentration and 4 μM protein, ΔCTC formed WT-like protofilaments, whereas ΔCTLC formed very few long protofilaments (Figure 8A). At high MgCl_2_ concentration and 4 μM protein, while ΔCTC protofilaments looked indistinguishable from WT, ΔCTLC formed large multifilament bundles similar to ΔCTL (Figure 8B, Figure 9A). This suggests that the GTPase domain is sufficient for polymerization and bundle formation, similar to observations for the GTPase domain of *B. subtilis* FtsZ (31). These observations also rule out the possibility that the ΔCTL bundles are the result of a difference in orientation of the CTC with respect to the polymerizing GTPase domain. We also observed similarly decreased apparent GTP hydrolysis rates for ΔCTLC and ΔCTC compared to ΔCTL and WT, respectively, suggesting that the effects of lacking the CTC on GTP hydrolysis and polymerization are independent of the effects of lacking the CTL (Figure 8C). At 2.5 mM MgCl_2_ and 4 μM protein concentrations, FtsZ, ΔCTL, ΔCTC and ΔCTLC showed GTP hydrolysis rates of 3.18 ± 0.16, 1.99 ± 0.09, 2.94 ± 0.19 and 1.50 ± 0.06 min^−1^ respectively. We conclude that the absence of the CTL is the primary contributor to the increased lateral interaction between protofilaments in ΔCTL and that there is no observable effect of the CTC on lateral interaction in this context.

**FIGURE 8:**
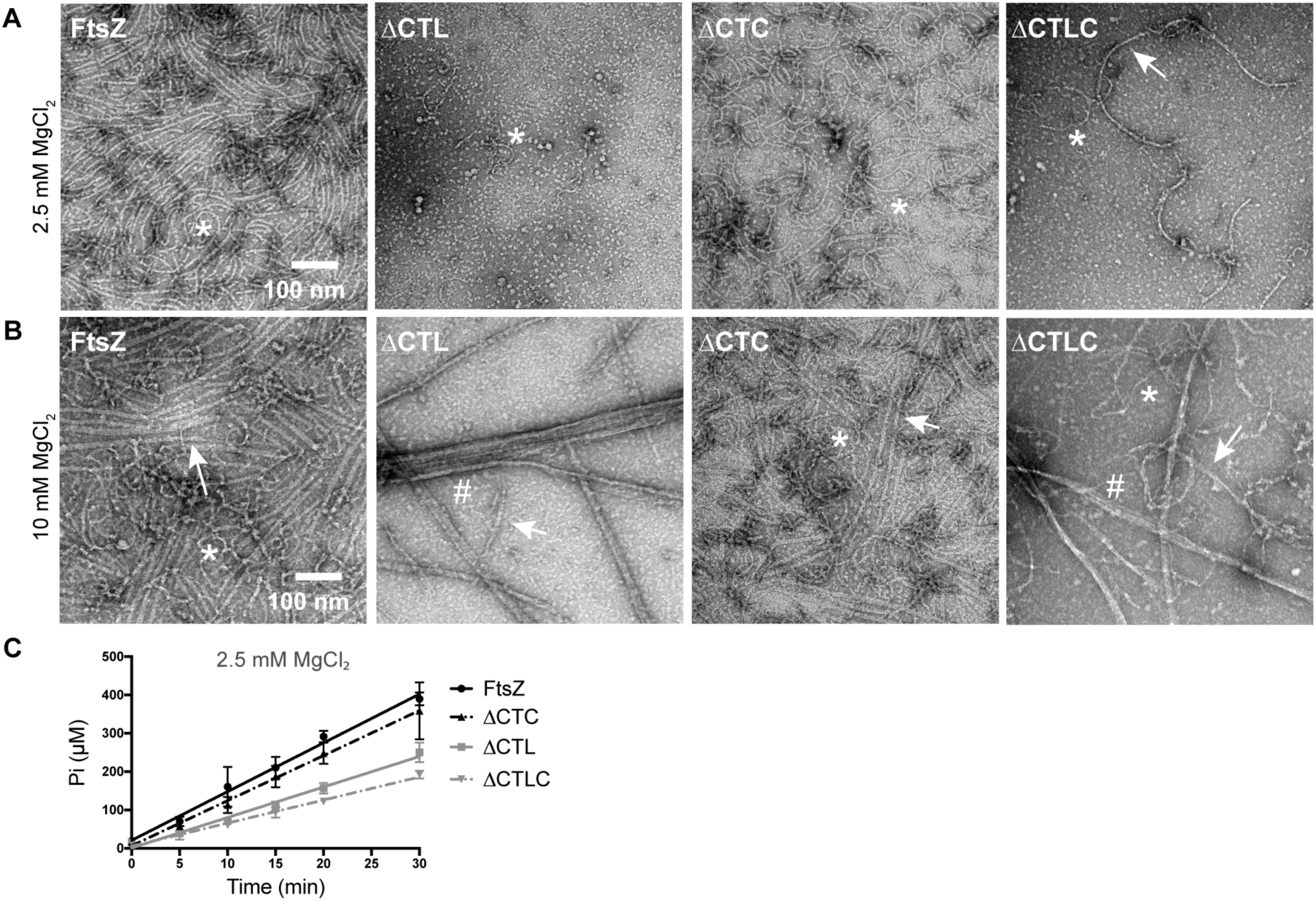
ΔCTL bundles form independent of the CTC. **A - B.** Electron micrographs of polymers formed by 4 μM FtsZ, ΔCTL, ΔCTC or ΔCTLC with 2 mM GTP and 2.5 mM MgCl_2_ **(A)** or 10 mM MgCl_2_ **(B)** spotted on grids 15 minutes after addition of nucleotide and stained with uranyl formate. Scale bar – 100 nm. * - single protofilaments, arrow – two or three filament bundles, # – multifilament bundles. **C.** Inorganic phosphate (P_i_) concentration in solution over time for reactions containing 4 μM FtsZ, ΔCTL, ΔCTC or ΔCTLC with 2 mM GTP and 2.5 mM MgCl_2_ (n = 3). Error bars represent standard deviation. Straight lines indicate linear fits of averages.

### FzlC binding reduces lateral interaction between ΔCTL protofilaments

FtsZ binding partners such as FzlA and ZapA have been implicated in increased lateral interaction (bundling) of protofilaments (12, 13, 32, 33). The increased bundling seen with ΔCTL appears more pronounced than the previously characterized effects of binding partners of *C. crescentus* FtsZ at similar polymerization conditions. Consequently, we investigated the influence of binding partners of FtsZ on the increased bundles observed for ΔCTL. While ZapA has been implicated in increased bundling of FtsZ in *E. coli*, no such effect has been observed for *C. crescentus* ZapA (30). WT FtsZ protofilaments look indistinguishable with or without ZapA (Supplementary figure 1C). We also failed to observe any difference in lateral interactions or structures of protofilaments formed by ΔCTL in the presence or absence of ZapA (Supplementary figure 1C).

Next, we characterized the effects of FzlC on protofilament structure (Figure 9). FzlC was recently identified as a membrane anchoring protein for FtsZ in *C. crescentus* (12, 24). *In vivo*, FzlC binds specifically to FtsZ’S CTC and recruits it to the membrane prior to the arrival of FtsA (12, 24). *In vitro*, His_6_-YFP-FzlC can recruit FtsZ-CFP to membranes of giant unilamellar vesicles in a CTC-dependent manner (24). At either low or high MgCl_2_ concentration, we did not observe any effect of His_6_-YFP-FzlC (2 μM) on protofilament structure for WT FtsZ (4 μM) (Figure 9B, FtsZ). Surprisingly, however, we found that the presence of His_6_-YFP-FzlC almost completely prevented bundling of ΔCTL at high MgCl_2_ concentration (Figure 9B, ΔCTL). In fact, the presence of His_6_-YFP-FzlC restores WT-like protofilament structures for ΔCTL. The presence of His_6_-YFP-FzlC did not affect the formation of multifilament bundles by ΔCTLC at high MgCl_2_ concentration, suggesting that FzlC’S ability to disrupt ΔCTL bundle formation requires its binding to the CTC (Figure 9B, ΔCTLC). His_6_-YFP-FzlC did not have an observable effect on ΔCTC, except for an overall reduction in the number of filaments on the grid (Figure 9B, ΔCTC). Thus, FzlC binding disrupts the increased lateral interaction in ΔCTL in a CTC-dependent manner.

**FIGURE 9:**
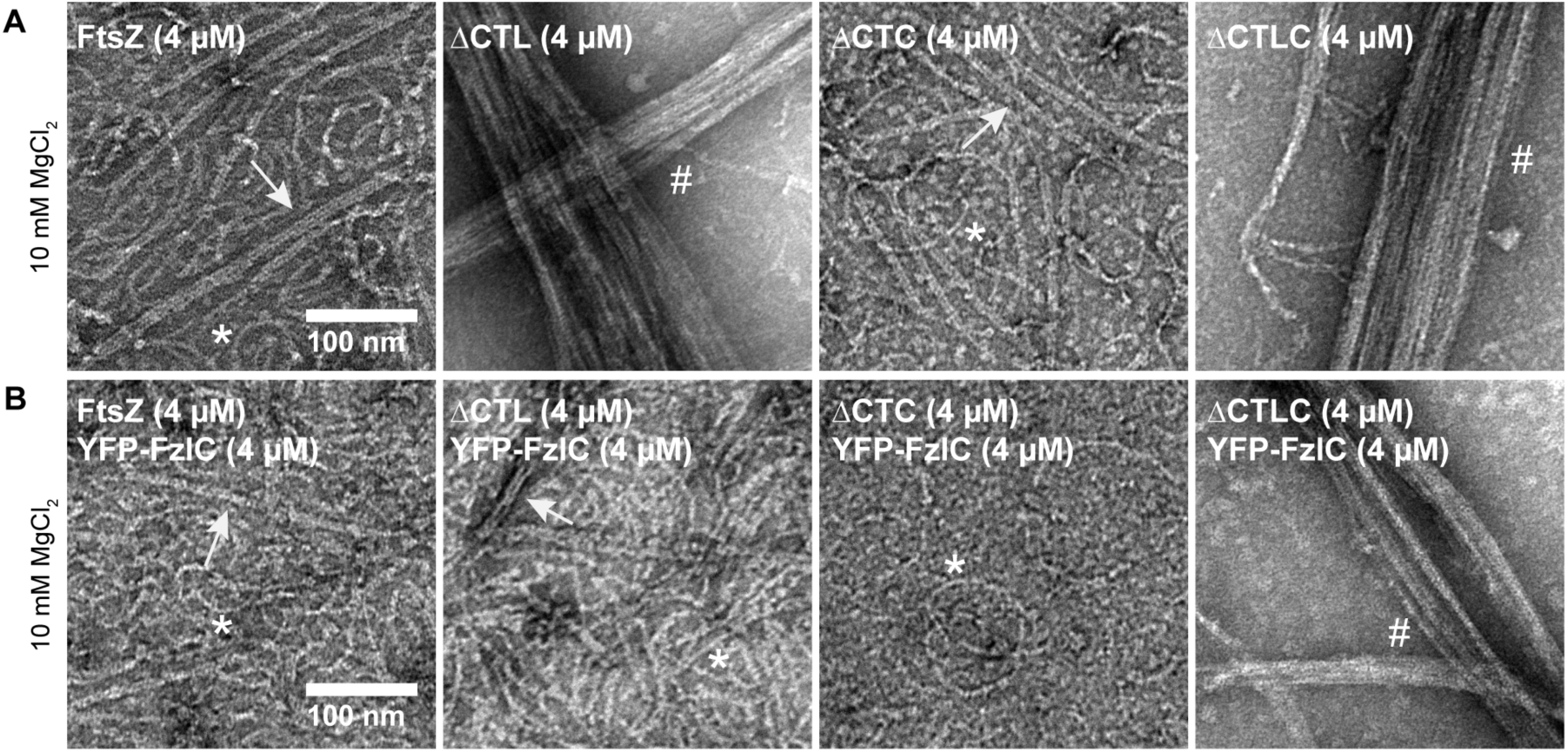
FzlC disrupts ΔCTL bundling in a CTC-dependent manner. **A - B.** Electron micrographs of polymers formed by 4 μM FtsZ, ΔCTL, ΔCTC or ΔCTLC with 2 mM GTP, 10 mM MgCl_2_ and 165 mM KCl in the absence **(A)** or presence **(B)** of 4 μM YFP-FzlC spotted on grids 15 minutes after addition of nucleotide. The difference in KCl concentration for this experiment is due to the high KCl concentration (300 mM) in FzlC storage buffer and low stock concentration of YFP-FzlC. Scale bars – 100 nm. * - single protofilaments, arrow – two or three filament bundles, # – multifilament bundles.

### The CTL influences FzlA-induced bundling

FtsZ protofilaments form helical bundles in the presence of FzlA *in vitro* (12). A recent characterization of FzlA from our lab confirmed that FzlA does not require the CTL or CTC to bind FtsZ^3^. We therefore assessed the effects of FzlA on protofilaments formed by CTL variants by TEM (Figure 10, refer to Supplementary figures 2, 3 for larger images). Our laboratory recently determined that, while WT FtsZ robustly forms helices with His_6_-tagged FzlA at pH 7.2 (12), it more robustly forms helices with untagged FzlA at pH 6.5^3^. We therefore first examined the structures formed by the CTL variants with untagged FzlA at lower pH. Under these conditions, we observed helical bundles only for WT FtsZ (Figure 10B, 10D, Supplementary figures 2B, 2D). However, we did observe an apparent stabilizing effect of FzlA on ΔCTL and L14 polymers (Figure 10A – 10D, Supplementary figure 2). At 2.5 mM MgCl_2_ concentration, we did not observe many protofilaments of ΔCTL or L14 in the absence of FzlA (Figure 10A, Supplementary figure 2A). However, in the presence of FzlA, we readily observed WT-like single filaments for both proteins (Figure 10B, Supplementary figure 2B). Moreover, we also observed straight double- and triple-filament bundles for ΔCTL with FzlA under these conditions (Figure 10B, Supplementary figure 2B). It is unclear if this is an effect on protofilament stability in solution or on binding EM grids. At 10 mM MgCl_2_ concentration, we observed thick bundles for ΔCTL that were unaffected by the presence of FzlA (Figure 10C, 10D, Supplementary figure 2C, 2D). Despite the increased number of protofilaments observed for L14 in the presence of FzlA, we still failed to observe very large bundles like those seen for ΔCTL either at low or high MgCl_2_ concentration (Figure 10B, 10D, Supplementary figure 2B, 2D). The structures formed by *Hn*CTL appeared similar in the presence or absence of untagged FzlA at pH 6.5 and either low or high magnesium concentration (Figure 10A – D, Supplementary figure 2).

**FIGURE 10:**
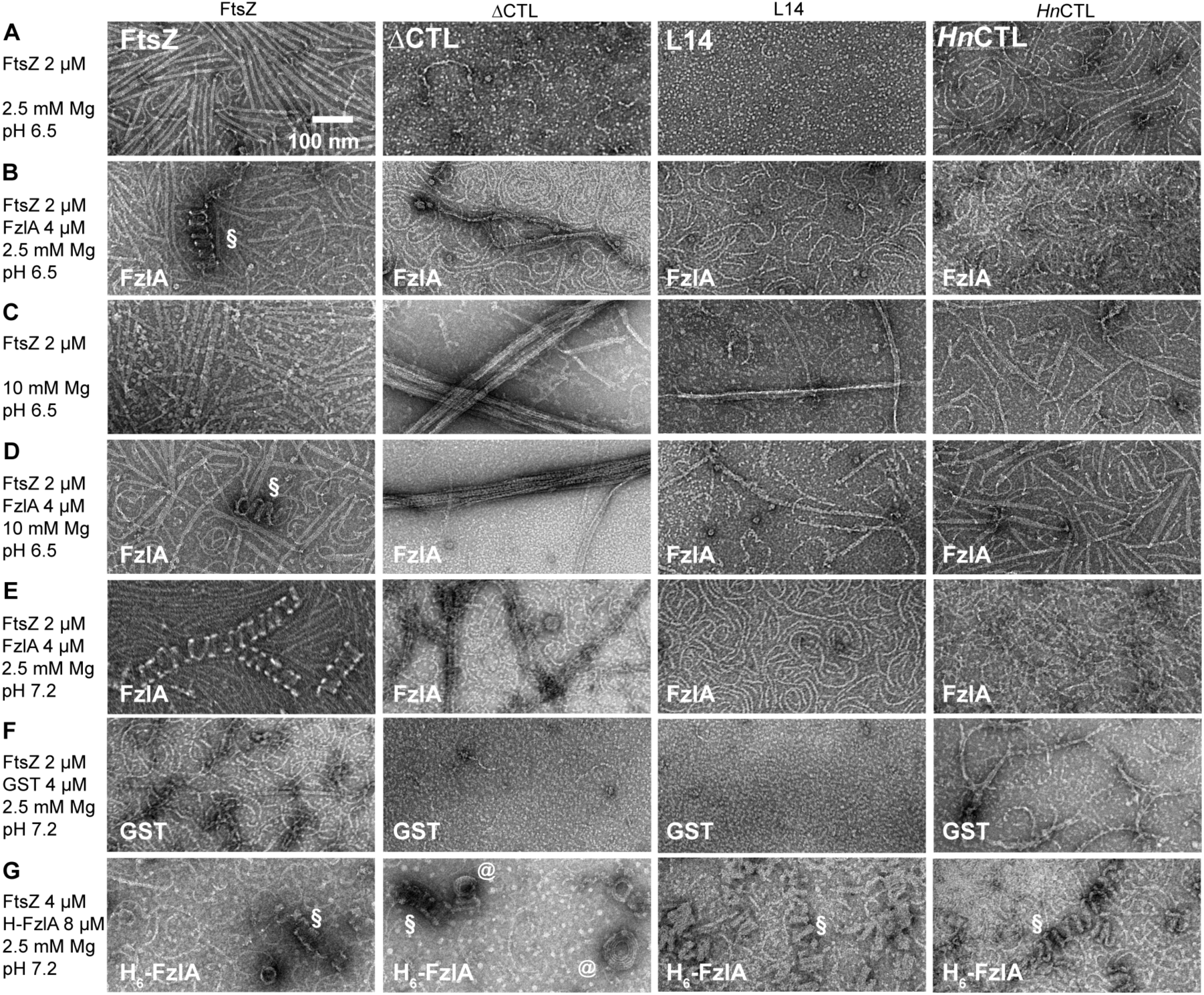
FzlA-induced bundling of FtsZ protofilaments into helices is CTL-dependent. **A - F.** Electron micrographs of polymers formed by 2 μM FtsZ CTL variants in the presence or absence of 2 μM FzlA at different pH (6.5 vs 7.2) with 2.5 mM or 10 mM MgCl_2_ concentration and 2 mM GTP spotted on grids 15 minutes after addition of nucleotide and stained with uranyl formate. **G.** Electron micrographs of polymers formed by 2 μM FtsZ CTL variants in the presence of 2 μM Gluthione S-transferase (GST) at pH 7.2 with 2.5 mM MgCl_2_ concentration and 2 mM GTP spotted on grids 15 minutes after addition of nucleotide and stained with uranyl formate. **H.** Electron micrographs of polymers formed by 4 μM FtsZ CTL variants in the presence of 8 μM His_6_-FzlA at pH 7.2 with 2.5 mM MgCl_2_ concentration and 2 mM GTP spotted on grids 15 minutes after addition of nucleotide and stained with uranyl formate. Scale bar – 100 nm. § - Helical bundles of FtsZ (or CTL variant) with FzlA, @ - spiral structures seen for ΔCTL with His_6_-FzlA.

We also assessed the effects of untagged FzlA on the structures of polymers formed by CTL variants at the presumably more physiological pH of 7.2 (Figure 10E, Supplementary figure 3). Again, we observed helical bundles only for WT FtsZ, and we observed a stabilizing effect of FzlA on ΔCTL and L14 (Figure 10E, Supplementary figure 3A, 3B). This effect was specific to FzlA since Glutathione S-transferase, which has structural homology to FzlA but does not bind FtsZ, did not affect polymer structure or density on grids under identical conditions (Figure 10F).

To use a more physiological pH while simultaneously favoring FzlA-induced bundling, we decided to use 8 μM His_6_-FzlA and 4 μM FtsZ or CTL variants at 2.5 mM MgCl_2_ concentration and pH 7.2. Under these conditions, all of the linker variants formed helical bundles similar to those observed for WT FtsZ (Figure 10G, Supplementary figure 3C). We observed fewer and shorter helical bundles for ΔCTL:His_6_-FzlA compared to the other CTL variants or WT. Surprisingly, we also observed flat spiral or coiled structures for ΔCTL:His_6_-FzlA (Figure 10G (denoted by @), Supplementary figure 3C inset). These structures resemble a highly-curved filament or filament bundle coiling onto itself. We hypothesize that these structures could result from increased self-interaction laterally between adjacent turns of helical double or triple filament bundles. Overall, our observations indicate that the CTL can affect FzlA-induced protofilament bundling and curvature. This is surprising since FzlA does not require the CTL for binding FtsZ^3^ and suggests that the CTL may influence the FzlA-FtsZ interaction through its intrinsic effects on the FtsZ assembly.

## DISCUSSION

In this study, we demonstrate a role for the intrinsically disordered C-terminal linker region of *C. crescentus* FtsZ in regulating polymer structure and dynamics *in vitro.* FtsZ variants lacking the CTL entirely (ΔCTL, ΔCTLC) exhibit increased propensity to bundle and reduced GTP hydrolysis rate and turnover compared to the other CTL variants or WT (Figure 8). The exceptionally long and thick straight bundles of ΔCTL protofilaments form independent of GTP hydrolysis rates, since we see long, thick bundles only for ΔCTL even with the slowly hydrolyzed GTP analog GMPCPP (Figure 2B, 2C, 3B, 3C). Combining biochemistry, spectroscopy and electron microscopy approaches, we confirmed that these bundles are not artifacts of observing a poorly polymerizing or depolymerizing FtsZ mutant under conditions that promote crowding (high MgCl_2_ or protein concentration). We confirmed that WT FtsZ and ΔCTL can copolymerize and observed that FtsZ:ΔCTL bundled copolymers appear much longer than FtsZ polymers (Figure 7). We show that FzlC binding to the CTC domain can disrupt the ability of ΔCTL to form bundles (Figure 9). This observation suggests a role for the membrane anchoring protein FzlC in regulating FtsZ polymer structure and dynamics in a CTC dependent manner. Additionally, it suggests that ΔCTL has the ability to form gently curved single filaments given the right conditions and/or binding partners, but intrinsically prefers to form bundles. Finally, we described a previously unidentified role for the CTL in facilitating FzlA to form helical bundles of FtsZ (Figure 10, Supplementary figures 2, 3). Overall, our characterization of the CTL variants and their interaction with FtsZ-binding proteins under different polymerizing conditions provide novel insights into the contributions of the CTL to FtsZ polymerization dynamics.

Based on our observations, we propose the following roles for the disordered linker in regulating lateral and longitudinal interactions between FtsZ monomers in protofilaments (Figure 11). Following addition of GTP, WT FtsZ undergoes fast polymerization (activation, nucleation, elongation); reaches a steady state characterized by dynamic turnover, annealing and fragmentation and transient lateral interactions; and eventually depolymerizes following exhaustion of GTP. The CTL likely plays a role in maintaining optimal interactions between protofilaments laterally, acting as flexible, charged “repulsive brushes” extending orthogonally around the protofilaments. Such a mechanism of regulating lateral interaction between polymers has been described for the intrinsically disordered C-terminal region of neurofilaments (34). Additionally, the CTL could influence longitudinal interactions between monomers through a similar electrostatic repulsion mechanism and thereby influence monomer on and off rates and/or fragmentation and annealing. In the case of ΔCTL, lateral interactions are more likely to form and less likely to break apart, with a strong propensity for the protofilaments to form long multifilament bundles. Moreover, ΔCTL might lack the repulsive interactions between adjacent monomers in a protofilament and therefore affect depolymerization and/or fragmentation. Together, these differences in lateral and longitudinal interactions would result in more stable polymers and slower turnover of monomers for ΔCTL.

**FIGURE 11:**
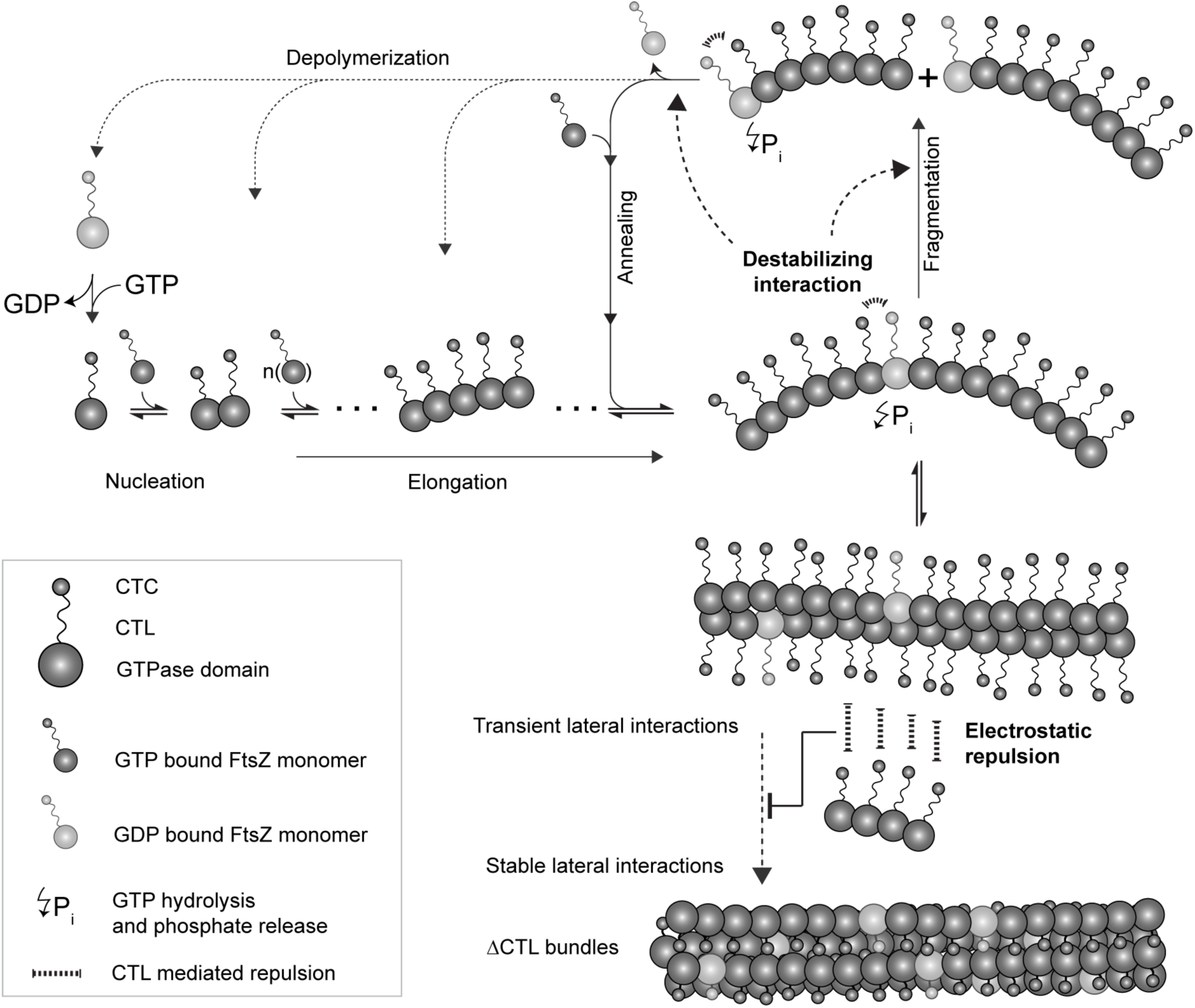
CTL regulates lateral interactions between FtsZ protofilaments and polymer turnover. Schematic illustrating polymerization dynamics of FtsZ and highlighting the proposed roles for the CTL in regulating longitudinal and lateral interactions. FtsZ monomers bind GTP, nucleate and assemble into short protofilaments that can elongate further and/or anneal together. GTP bound monomers within the protofilaments undergo GTP hydrolysis leading to a conformational change that results in destabilization of longitudinal interactions. The CTL may play a role in regulating monomer-monomer interactions that contribute to this destabilization, which leads to fragmentation and depolymerization. At steady state, the forward reactions – nucleation, elongation and annealing, balance out the reverse reactions, depolymerization and fragmentation, as long as GTP bound FtsZ monomers are available. FtsZ protofilaments can also form lateral interactions that are most likely disrupted prior to fragmentation and/or depolymerization. We propose that the CTL plays a role in regulating lateral interactions by functioning as a repulsive brush around FtsZ protofilaments. In the absence of the CTL, lateral interaction between protofilaments is stronger, thereby favoring the assembly of protofilaments into stable bundles. Moreover, in the absence of the CTL, protofilaments may be stabilized against fragmentation/depolymerization both by bundle formation and by direct effects of the CTL on longitudinal interactions, leading to formation of very long filaments.

We consistently observed reduced lateral interaction for *Hn*CTL (165 aa CTL, Figure 1A, Figure 2, 3, 10) protofilaments compared to WT (172 aa CTL, Figure 1A), by EM, pelleting, and light scatter assays. It is interesting to note that while *C. crescentus* CTL has a net charge of – 9 and pI of 4.66, *Hn*CTL has a net charge of – 18 and pI of 4.2. The reduction in lateral interaction for *Hn*CTL, which has a more negatively charged CTL than WT, further supports our model that electrostatic repulsion between protofilaments mediated by the CTL is important for optimal lateral interaction and bundle formation. A similar observation has been reported earlier for *B. subtilis* FtsZ CTL variants – replacing the CTL (50 aa) of BsFtsZ with CTL from *E. coli* (50 aa) or *A. tumefaciens* (50 aa) causes reduction in lateral interaction both by EM and light scatter (25). Whether the CTL sequence-dependent difference in protofilament bundling is due to CTL-CTL interactions or CTL-GTPase domain interactions between adjacent protofilaments needs further investigation.

Across species of bacteria, many members of the division machinery, such as FzlA and the Zap family of proteins, have been implicated in promoting interactions between FtsZ protofilaments *in vitro* and/or regulating the density of protofilaments in the Z-ring (12, 13, 32, 33, 35, 36). Genetic manipulations of these proteins result in cell division defects, suggesting a role for regulating interfilament interactions in the cytokinetic function of FtsZ (13, 30)^3^. At least for the formation of helical bundles *in vitro*, untagged FzlA appears to require the CTL (Figure 10). We have no evidence that FzlA interacts with the CTL. Thus, it is possible that the different intrinsic polymerization properties or lateral interactions of different CTL variants affects the curvature or polymer dynamics required for forming helical bundles upon FzlA binding. The His_6_-tagged variant of FzlA can form robust helices with all linker variants (Figure 10G), which might be due to altered interactions with FtsZ. Interestingly, His_6_-FzlA also causes ΔCTL to form unique spiral or coiled structures that have never been observed for WT FtsZ with FzlA again suggesting that His_6_-FzlA might have non-canonical interactions with FtsZ.

Many proteins that bind the CTC of FtsZ have been shown to affect Z-ring structure *in vivo* and/or affect FtsZ polymer structure *in vitro* (24, 37–39). It is unclear if and how these proteins regulate polymerization of FtsZ without interacting directly with the GTPase domain, since the CTC and the GTPase domain are linked by a long unstructured CTL. In the context of ΔCTL, the CTC does not obviously contribute to the regulation of protofilament structure or lateral interaction in *C. crescentus* FtsZ. However, FzlC binding to the CTC disrupts ΔCTL bundling. Whether FzlC is relevant for regulating interprotofilament spacing in WT FtsZ is unclear. It is curious, however, that while ZapA and FzlA, which bind the GTPase domain of FtsZ (13, 33)^3^, do not affect bundling of ΔCTL protofilaments, binding of FzlC to the CTC region disrupts bundling. It is possible that the binding of FzlC to the CTC of ΔCTL disrupts inter-protofilament lateral interactions at the GTPase domain by restricting the orientation of protofilaments. Alternatively, FzlC bound to the CTC of ΔCTL may perform a repulsive function to limit lateral interaction similar to the mechanism we propose for the CTL, itself. Interestingly, the effect of FzlC on preventing ΔCTL bundles occurs in the absence of membrane, indicating that this ability of FzlC, if relevant *in vivo*, is independent from its function as a membrane anchor for FtsZ. Further *in vitro* studies on the interaction between FzlC and FtsZ would be required to understand the mechanism underlying this observation and to determine if it extends to other factors that bind the CTC, such as FtsA. Differences in the effects of membrane anchoring proteins such as FzlC or FtsA on FtsZ polymerization could contribute to the roles of these proteins in cell division, in addition to their abilities to recruit FtsZ protofilaments to the membrane.

How transient are the bundles observed for ΔCTL? Does WT FtsZ also form similar bundles that dissociate more quickly and are difficult to capture for TEM? Our data from experiments using GMPCPP suggests that reducing turnover of FtsZ does not result in the same extent of bundling as observed for ΔCTL. While TEM provides very high spatial resolution of protofilaments and polymer structure, it has two major limitations. First, we cannot observe changes in structures over time. Second, differences in staining and affinity for the carbon-coated grids could bias the observer towards large bundles and/or well spaced polymers. To overcome these limitations and resolve FtsZ polymerization dynamics, we would require highresolution time-lapse microscopy and/or spectroscopy techniques.

Our characterization of the CTL variants and FtsZ mutants provides biochemical tools for resolving the roles of longitudinal and lateral interactions in regulating protofilament structure *in vitro* and *in vivo.* Similar studies of polymerizing proteins such as actin and microtubules in eukaryotes have potentiated a relatively thorough understanding of the regulation and function of their assembly properties. Our recent understanding that the Z-ring is more than a passive scaffold for the recruitment of cell wall enzymes and the role for its dynamics in regulating cytokinesis emphasizes the need for understanding its polymerization properties in molecular detail. Additionally, while our observations strengthen the correlation between polymerization dynamics *in vitro* and Z-ring structure and function *in vivo*, whether this correlation is suggestive of causation or the result of a confounding effect of the CTL on unrelated processes requires further investigation.

## EXPERIMENTAL PROCEDURES

### Purification of proteins

FtsZ and FtsZ variants (Figure 1A) including FtsZ_L72w_ and ΔCTL_L72W_ tryptophan mutants were purified using the protocol described previously (28). Briefly, pET21 vectors (pMT219 - FtsZ, pEG681 - ΔCTL, pEG723 – L14, pEG676 – *Hn*CTL, pEG765 - ΔCTC, pEG678 – ΔCTLC, pEG948 – FtsZ_L72w_ (29), pEG1077 – ΔCTL_L72W_) were used to express *ftsZ* or *ftsZ* variants in *E. coli* Rosetta cells induced with 0.5 mM IPTG at 37°C for 3 hours when OD_600_ reached 1.0. Cells were pelleted, resuspended in lysis buffer (50 mM Tris-HCl pH 8.0, 50 mM KCl, 1 mM EDTA, 10% glycerol, DNase I, 1 mM β-mercaptoethanol, 2 mM PMSF with cOmplete mini, EDTA-free Protease inhibitor tablet (Roche)), and lysed using lysozyme treatment (1 mg/mL) followed by sonication. After anion exchange chromatography (HiTrap Q HP 5 mL, GE Life Sciences) and elution with a linear gradient of KCl, the fractions containing the FtsZ variant were pooled and subjected to ammonium sulfate precipitation. The ammonium sulfate precipitates (at 20-35% saturation, depending on the variant) were verified for each FtsZ variant by
electrophoresis (SDS-PAGE) and Coomassie staining. The precipitate was resuspended in FtsZ storage buffer (50 mM HEPES-KOH pH 7.2, 50 mM KCl, 0.1 mM EDTA, 1 mM β-mercaptoethanol, 10% glycerol) and purified further using size-exclusion chromatography (Superdex 200 10/300 GL, GE Life Sciences), snap frozen in liquid nitrogen, and stored at −80°C in FtsZ storage buffer.

ZapA was purified as described previously (30). His_6_-SUMO-ZapA was produced from plasmid pEG620 in *E. coli* Rosetta cells induced with 0.5 mM IPTG for 4 hours at 37°C. Cells were pelleted, resuspended in ZapA lysis buffer (50 mM Tris-HCl pH 8.0, 100 mM NaCl, 20 mM imidazole, 10% glycerol) and lysed with lysozyme treatment (1 mg/ml lysozyme, 2.5 mg/ml MgCl_2_, 1 mM CaCl_2_ and 2 units/ml DNaseI) and sonication. His_6_-SUMO-ZapA was purified using a HisTrap FF 1 ml column (GE Life Sciences) and eluted with 300 mM imidazole. The His_6_-SUMO tag was cleaved overnight at 4°C during dialysis into ZapA lysis buffer using SUMO protease (His_6_-Ulp1) at a 100-fold molar excess. Cleaved His_6_-SUMO, uncleaved His_6_-SUMO-ZapA, and His_6_-Ulp1 were separated from untagged ZapA by passage over a HisTrap FF 1 ml column once again, this time collecting the unbound fraction. ZapA was further purified using anion exchange (HiTrap Q HP 1 mL, GE Life Sciences) in ZapA QA Buffer (50 mM Tris-HCl pH 8.0, 100 mM NaCl, 10% glycerol) with a linear gradient of NaCl, and was dialyzed into FtsZ storage buffer before snap freezing and storage at −80°C.

His_6_-YFP-FzlC was purified as described previously (24). Briefly, His_6_-YFP-FzlC was produced in *E. coli* Rosetta cells from plasmid pEG420 using 30 μM IPTG to induce expression overnight at 15°C. Cells were pelleted and resuspended in FzlC lysis buffer (50 mM Tris-HCl pH 8.0, 1 M KCl, 20 mM imidazole, 1 mM β-mercaptoethanol, and 20% glycerol, 2 mM PMSF with cOmplete mini, EDTA-free Protease inhibitor tablet (Roche)) and lysed using lysozyme treatment (with 1 mg/ml lysozyme, 2.5 mg/ml MgCl_2_, and 2 units/ml DNaseI) and sonication. Lysate was supplemented with 3 mM ATP (to reduce DnaK co-purification) and His_6_-YFP-FzlC was purified using affinity chromatography (His-Trap FF 1 ml column, GE Life Sciences) followed by gel filtration (Superdex 200 10/300 GL column, GE Life Sciences) and was stored in FzlC storage buffer (50 mM Tris-HCl pH 8.0, 300 mM KCl, 0.1 mM EDTA, 1 mM β-mercaptoethanol and 10% glycerol) at −80°C.

His_6_-FzlA was purified essentially as described previously (12). His_6_-FzlA was expressed from plasmid pEG327 in *E. coli* Rosetta cells by induction with 0.5 mM IPTG for 4 h at 30°C when OD_600_ reached 0.5. Cells were pelleted, resuspended in lysis buffer (50 mM Tris-HCl pH 8.0, 300 mM NaCl, 20 mM imidazole, 10% glycerol, 2 mM PMSF with cOmplete mini, EDTA-free Protease inhibitor tablet (Roche)), and lysed using lysozyme treatment (1 mg/mL lysozyme, 2 units/mL DNAse I, 2.5 mM MgCl2) and sonication. His_6_-FzlA protein was isolated using affinity chromatography (HisTrap FF 1 ml, GE Life Sciences), eluted with 300 mM imidazole, and dialyzed into FzlA storage buffer (50 mM HEPES-KOH pH 7.2, 300 mM KCl, pH 8.0, 10% glycerol).

For the purification of untagged FzlA, His_6_-SUMO-FzlA was expressed from plasmid pEG994 in *E. coli* Rosetta cells by induction with 0.5 mM IPTG for 4 h at 30°C. His_6_-SUMO-FzlA was purified and cleaved to FzlA using a similar protocol to ZapA purification from His_6_-SUMO-ZapA, with a few changes - FzlA lysis buffer (50 mM HEPES-KOH pH 7.2, 300 mM KCl, 20 mM imidazole, 10% glycerol) was used for lysis. Affinity chromatography (HisTrap FF 1ml column, GE Life Sciences) followed by SUMO protease cleavage produced untagged FzlA that was further purified as the unbound fraction of another passage of a HisTrap FF 1ml column (GE Life Sciences). Untagged FzlA was dialyzed and stored in 50 mM HEPES-KOH pH 7.2, 300 mM KCl, 10% glycerol.

GST was produced from pGEX4T1 in *E. coli* Rosetta cells grown at 30°C and induced at OD_600_ of 0.5 with 0.5 mM IPTG for 3 h. Cells were resuspended in phosphate buffered saline (PBS) with 300 mM KCl, lysed with 1 mg/mL lysozyme treatment and sonication, and purified by affinity chromatography (Glutathione Sepharase 4B column, GE Healthcare). Protein was eluted with 10 mM glutathione and dialyzed into GST storage buffer (50 mM HEPES-KOH pH 7.2, 50 mM KCl, 0.1 mM EDTA) before snap freezing and storage at −80°C.

His_6_-FzlA, FzlA and GST were buffer exchanged into FtsZ storage buffer (50 mM HEPES-KOH pH 7.2, 0.1 mM EDTA, 50 mM KCl, 10% glycerol), prior to addition to polymerization reactions with FtsZ.

### Polymerization buffer and conditions

HEK50 pH 7.2 buffer (50 mM HEPES-KOH pH 7.2, 0.1 mM EDTA, 50 mM KCl) was used for FtsZ polymerization assays, unless specified otherwise. 2.5 mM MgCl_2_ (low magnesium) or 10 mM MgCl_2_ (high magnesium) were used as mentioned in results and figure legends. GTP or GMPCPP were used at 2 mM or 0.2 mM concentrations, respectively, unless specified otherwise. For experiments with FtsZ-CTL variants and FzlA at pH 6.5, MESK50 pH 6.5 buffer (50 mM MES-KOH pH 6.5, 50 mM KCl) was used. All reactions were carried out at room temperature. When additional proteins were added, the same volume of the corresponding storage buffer was added to a corresponding control reaction with no added protein.

### GTP hydrolysis rate measurement

GTP hydrolysis rates were measured using malachite green dye based reporter for inorganic phosphate as described previously (28). At the beginning of the reaction, 2 mM GTP was added to FtsZ or FtsZ variants in polymerization buffer at 2.5 or 10 mM MgCl_2_ concentration. Reaction was stopped using quench buffer (50 mM HEPES-KOH pH 7.2, 21.3 mM EDTA, 50 mM KCl) and inorganic phosphate in the solution was measured using SensoLyte MG Phosphate Assay Kit Colorimetric (AnaSpec, Inc, Fremont, California).

### Polymerization kinetic assays

FtsZ polymerization was measured using Fluoromax-3 spectrofluorometer (Jobin Yvon Inc.) to measure right angle light scatter (excitation and emission at 350 nm, 2 nm slits). Tryptophan fluorescence experiments using FtsZ_L72W_ and ΔCTL_L72W_ mutants were performed using the same equipment, with excitation and emission at 295 nm and 344 nm respectively with 2 nm slits. Since GTP is fluorescent at these excitation/emission conditions, 50 μM GTP was used for the tryptophan assay (29). Similarly, 100 μM GMPCPP was used for the assay with increasing MgCl_2_ concentration (Figure 6). Note that addition of 100 μM GMPCPP does not cause significant increase in fluorescence signal at 344 nm, by itself (Figure 6). In both right angle light scatter and tryptophan fluorescence experiments, measurements were taken every 10 seconds.

### High speed pelleting assay

Steady state polymer mass was measured using high speed pelleting assay as described previously (12, 28). Briefly, FtsZ or FtsZ variants in storage buffer were centrifuged at 250,000 × g for 15 minutes at 4 °C to pellet and remove non-specific aggregates in the absence of GTP. The clarified FtsZ variant (from the supernatant) was then incubated for 15 minutes with 2 mM GTP and 10 mM MgCl_2_ in HEK50 pH 7.2 buffer with 0.05% Triton X-100. Polymers were then pelleted by ultracentrifugation at 250,000 × g for 15 minutes at 25 °C. The amount of protein in the supernatant and pellet were determined using SDS-PAGE and Coomassie staining followed by densitometry using ImageLab (Bio-Rad Laboratories, inc., USA).

### Transmission electron microscopy

Polymers formed by FtsZ or FtsZ variants (in the presence or absence of FtsZ binding proteins) were visualized by TEM. Reaction mixtures of the relevant proteins in FtsZ polymerization buffer were spotted onto glow-discharged carbon coated copper grids (Electron Microscopy Sciences, Hatfield, PA) at least 15 minutes after addition of GTP except when noted otherwise, blotted and stained twice with 0.75 % uranyl formate for 2 minutes. The grids were dried and imaged using a Philips/FEI BioTwin CM120 TEM (operated at 80 kV) equipped with an AMT XR80 8 megapixel CCD camera (AMT Imaging, USA).

## Acknowledgements

We would like to thank Harold Erickson (Duke University) for providing the construct for the purification of FtsZ_L72W_. We also thank the members of Goley lab: Elizabeth Meier, PJ Lariviere, and Selam Woldemeskel, for guidance with optimizing assays with FzlC, FzlA, and ZapA respectively; Allison Daitch and Chris Mahone for useful discussions about the manuscript. We would also like to thank Mike Delannoy, Barbara Smith, and the Microscopy Facility of Johns Hopkins School of Medicine for assistance with the electron microscopy imaging. Finally, we thank Jie Xiao and Anthony Vechiarelli for providing useful insights into FtsZ polymerization.

## Author contributions

KS and EDG designed the study. KS performed the experiments. KS and EDG analyzed the results, wrote the paper and approved the final version of the manuscript.

## Footnotes

1 This work was supported by the National Institutes of Health grant R01GM108640 awarded to EDG.

2 The abbreviations used are: FtsZ – Filamentous Temperature Sensitive mutant Z, CTL – C-Terminal Linker, CTC – C-terminal conserved peptide, WT – Wildtype, FzlC – FtsZ linked protein C, FzlA – FtsZ linked protein A, ZapA – Z-ring associated protein A, MgCl_2_ – Magnesium chloride, TEM – Transmission Electron Microscopy, GMPCPP – Guanosine-5’-[(α,β)-methyleno]triphosphate, KCl – Potassium chloride, aa – amino acids, IPTG – Isopropyl β-D-1-thiogalactopyranoside.

3 Patrick J Lariviere, Piotr Szwedziak, Jan Löwe, and Erin D Goley, manuscript in revision.

